# Structural Determinants of *Vibrio cholerae* FeoB Nucleotide Promiscuity

**DOI:** 10.1101/2024.05.22.595361

**Authors:** Mark Lee, Kate Magante, Camilo Gómez-Garzón, Shelley M. Payne, Aaron T. Smith

## Abstract

Ferrous iron (Fe^2+^) is required for the growth and virulence of many pathogenic bacteria, including *Vibrio cholerae* (*Vc*), the causative agent of the disease cholera. For this bacterium, Feo is the primary system that transports Fe^2+^ into the cytosol. FeoB, the main component of this system, is regulated by a soluble cytosolic domain termed NFeoB. Recent reanalysis has shown that NFeoBs can be classified as either GTP-specific or NTP-promiscuous, but the structural and mechanistic bases for these differences were not known. To explore this intriguing property of FeoB, we solved the X-ray crystal structures of *Vc*NFeoB in both the apo and GDP-bound forms. Surprisingly, this promiscuous NTPase displayed a canonical NFeoB G-protein fold like GTP-specific NFeoBs. Using structural bioinformatics, we hypothesized that residues surrounding the nucleobase could be important for both nucleotide affinity and specificity. We then solved the X-ray crystal structures of N150T *Vc*NFeoB in the apo and GDP-bound forms to reveal H-bonding differences surround the guanine nucleobase. Interestingly, isothermal titration calorimetry revealed similar binding thermodynamics of the WT and N150T proteins to guanine nucleotides, while the behavior in the presence of adenine nucleotides was dramatically different. AlphaFold models of *Vc*NFeoB in the presence of ADP and ATP showed important conformational changes that contribute to nucleotide specificity among FeoBs. Combined, these results provide a structural framework for understanding FeoB nucleotide promiscuity, which could be an adaptive measure utilized by pathogens to ensure adequate levels of intracellular iron across multiple metabolic landscapes.

## INTRODUCTION

Iron (Fe) is an essential nutrient for nearly all lifeforms due to its use as a cofactor in numerous biochemical processes, including oxidative phosphorylation, *de novo* DNA synthesis, and nitrogen fixation, among others(1-5). To harness the power of this element, iron must first be acquired from the environment before it can be biologically incorporated into proteins and enzymes. For many organisms, including most bacteria, the prevalent environmental oxidation state of iron dictates its mode of acquisition. For example, the highly insoluble ferric iron (Fe^3+^) is prevalent in oxic environments, and bacteria will commonly deploy siderophores to solubilize and to capture this form of iron. Membrane receptors then translocate the siderophore-chelated iron into the cytosol, where the iron is either removed by reductive dissociation or by cleavage of the siderophore(6, 7). Additionally, some pathogenic bacteria can sequester either free heme (iron protoporphyrin IX) or utilize hemophores to remove heme from host hemoproteins (*e*.*g*., hemoglobin and myoglobin). Like siderophore-mediated uptake, membrane receptors then translocate the heme into the cytosol where the heme is either recycled or destroyed to remove the iron contained within(8-11). In contrast, when living within anoxic or acidic environments, bacteria commonly encounter the more labile, but also more reactive, ferrous iron (Fe^2+^)(12-15). Unfortunately, despite its strong contribution to bacterial metal homeostasis and pathogenesis, the mechanisms of bacterial Fe^2+^ acquisition are poorly understood compared to the mechanisms of Fe^3+^ uptake and heme acquisition.

While some auxiliary Fe^2+^ transport systems have been identified, the <u>fe</u>rrous ir<u>o</u>n transport (Feo) system is the most widely distributed and conserved Fe^2+^ acquisition system across the prokaryotic domain(12-15), although Feo’s precise mechanism of function remains unclear. Canonically, the *feo* operon encodes for three proteins, FeoA, FeoB, and FeoC(16), although FeoC is the least conserved of these proteins(13-15, 17) (Fig. 1). FeoA and FeoC are known to be small (*ca*. 7-10 kDa) cytosolic proteins, while FeoB is a large (*ca*. 80-100 kDa) polytopic transmembrane protein that contains an N-terminal soluble G-protein-like domain termed NFeoB(12, 14, 18). The roles of FeoA and FeoC remain somewhat enigmatic; however, these proteins have been shown to interact with NFeoB *in vitro(19)*, FeoA appears to regulate GTP hydrolysis *in vitro(17)*, and some FeoCs bind oxygen-sensitive [Fe-S] clusters, presumably for regulatory purposes(20). *In vivo*, several observations indicate that both proteins interact with FeoB and are required for Feo-dependent iron uptake in *Vibrio cholerae* (*Vc*)(21, 22), the pathogenic bacterium responsible for the diarrheal disease cholera. Interestingly, bacterial two hybrid (BACTH) systems demonstrated that both *Vc*FeoA and *Vc*FeoC were found to interact with intact *Vc*FeoB in the cell(23), although the precise nature of this complex and its mechanism are still unclear.

**Fig. 1.**
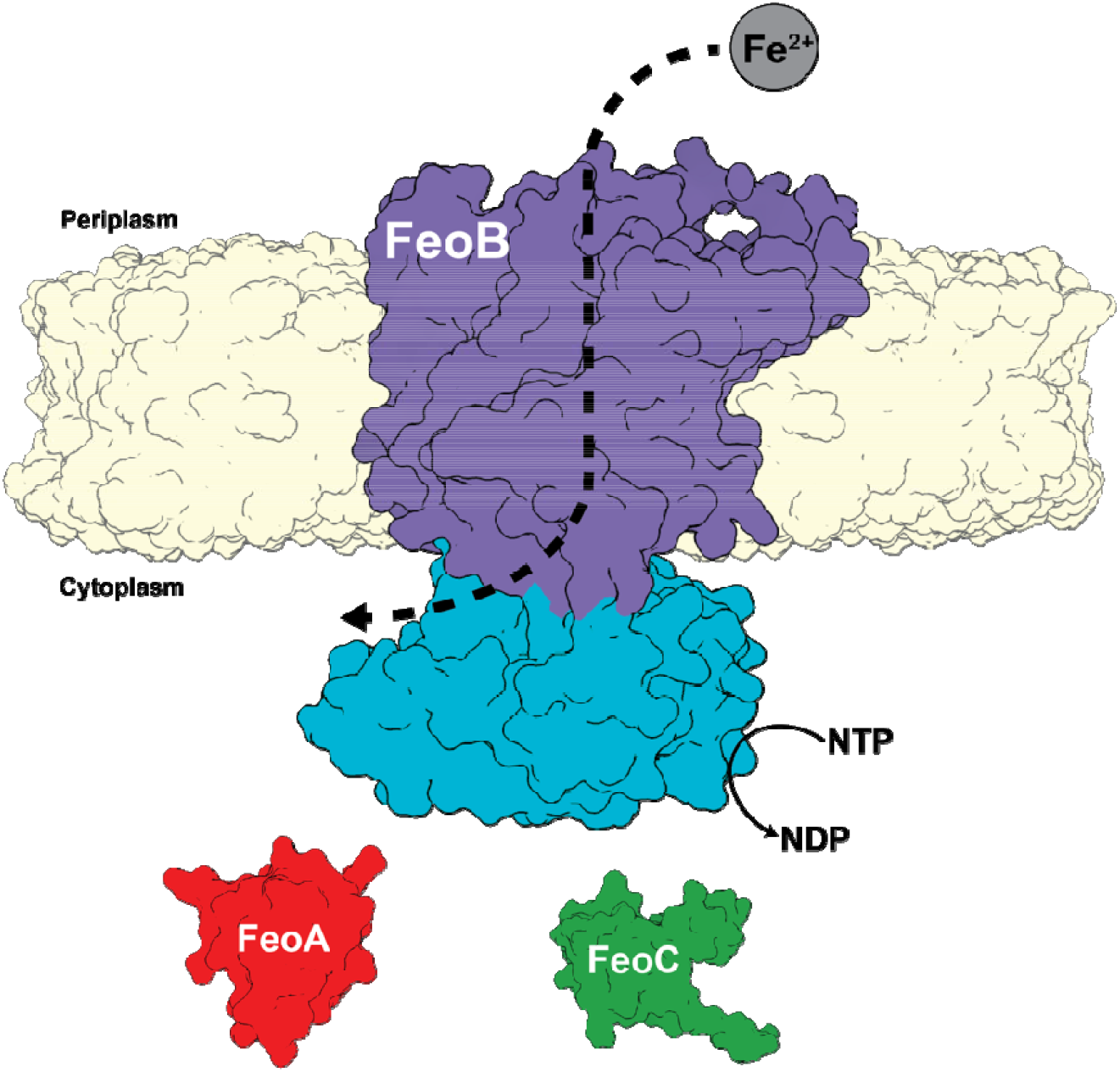
Cartoon depiction of the tripartite *Vibrio cholerae* ferrous iron transport (Feo) system. In *V. cholerae* the Feo system consists of three proteins: FeoA (colored in red) and FeoC (colored in green), both of which are cytosolic, and a polytopic transmembrane protein, FeoB (colored in purple), with soluble N-terminal domain termed NFeoB (colored in teal). The NFeoB domain of *V. cholerae* FeoB was recently discovered to be nucleotide promiscuous and is best classified as an NTPase rather than a strict GTPase.

FeoB from *V. cholerae* was recently shown to hydrolyze ATP, GTP, and other NTPs *in vitro*, and this function could serve to supply *V. cholerae* with Fe^2+^ *in vivo*, indicating that this FeoB may be better classified as an NTPase rather than a strict GTPase(24, 25). This promiscuity for NTP consumption was then shown to occur *in vitro* for other FeoBs from a handful of infectious bacterial species such as *Helicobacter pylori, Streptococcus mutans, Staphylococcus aureus*, and *Bacillus cereus*, which could suggest that NTP promiscuity may be a common theme used by some pathogenic bacteria to acquire iron and to establish infection. NFeoB proteins contain generally conserved G-motifs that are common amongst G-proteins and are responsible for binding to different segments of the guanine nucleotide, with G1, G2, and G3 binding to the α-, β-, and γ-phosphates, while the G4 and G5 motifs interact with the nucleobase(26). Sequence analyses suggested that differentially conserved residues within the G4 and G5 motifs might be important for guanine recognition and for NTPase activity in FeoB(27, 28). Specifically, the G5 motif residues Ser148 and Asn150, when altered, displayed significant effects on ATPase activity, but minimal effects on GTPase activity, indicating that these G5 residues may play a critical role in nucleotide discrimination(24). However, the structural basis of nucleotide promiscuity in FeoB remained unknown, precluding a more comprehensive understanding of this unique aspect of the Feo system.

In this work, we have structurally and biophysically characterized the *Vc*NFeoB domain in order to understand its nucleotide promiscuity. Using X-ray crystallography, we determined the structures of apo and GDP-bound *Vc*NFeoB, which reveals a conserved NFeoB fold composed of a G-protein like domain tethered to a GDI domain, despite the nucleotide promiscuous nature of *Vc*FeoB. Additionally, we determined the X-ray crystal structures of apo and GDP-bound N150T *Vc*NFeoB, and we show how differences in residues at the G5 motif alter the hydrogen-bonding interactions surrounding the guanine nucleobase. Isothermal titration calorimetry (ITC) was used to determine substrate affinities and stoichiometries of different nucleotides to both the wild-type and variant *Vc*NFeoBs. Intriguingly, we demonstrate dramatic differences in the behavior of the *Vc*NFeoB towards GDP/GMP-PNP binding compared to ADP/AMP-PNP binding, which could be rationalized using AlphaFold modeling. Taken together, these findings provide a structural framework for understanding the nucleotide promiscuity of *Vc*FeoB, which could be leveraged for future developments of targeted therapeutics to tackle issues of *V. cholerae* pathogenesis, as recently demonstrated(29).

## RESULTS

### The VcNFeoB NTPase Domain Purifies as a Nucleotide-Free Monomer that Displays Broad NTPase Activity

To prepare the *Vc*NFeoB NTPase domain for crystallization trials, we overproduced in *E. coli* a previously designed construct that encodes for a non-cleavable (His)_6_-tagged version of the protein (*i*.*e*., *Vc*NFeoB(His)_6_))(24). As this protein was not initially suitable for crystallization from immobilized metal-affinity chromatography (IMAC), we added two additional purification steps: anion exchange chromatography (AEX) and size-exclusion chromatography (SEC), both of which revealed interesting biophysical properties of *Vc*NFeoB(His)_6_. First, regarding AEX, we monitored the 260 nm / 280 nm ratio through the entire chromatography process, and noted a normal value of ≈0.6, indicating that *Vc*NFeoB(His)_6_ does not co-purify with nucleotide, unlike previous reports of *E. coli* NFeoB overproduced in *E. coli(30)*. Second, SEC of either crudely purified *Vc*NFeoB(His)_6_ (IMAC only) or polished *Vc*NFeoB(His)_6_ (after SEC) showed only the presence of a dominant monomeric species in solution (Fig. S1), consistent with our previous *in vitro* studies on Feo proteins (from *V. cholerae* and others) recombinantly produced in *E. coli*. It is possible that FeoA and/or FeoC may be necessary to induce FeoB oligomerization, and *in vivo* studies have suggested this to be the case for *Vibrio cholerae(23)*. However, other Feo systems appear to be functional monomers *in vitro*, and this highly pure, monomeric *Vc*NFeoB(His)_6_ domain was active against multiple nucleotide triphosphates, most notably ATP and GTP, as previously described(24), thus the precise oligomeric state of FeoB remains controversial and requires further exploration. For all subsequent constructs (*vide infra*), a similar overproduction and purification process was employed, producing highly pure, homogeneous, and monomeric protein (Fig. S1).

### The Structure of the Apo VcNFeoB NTPase Domain Reveals a Typical NFeoB Fold

In order to characterize the three dimensional structure of the *Vc*NFeoB NTPase domain, we sought to crystallize the *Vc*NFeoB(His)_6_ protein. Crystals of the apo domain were generated and ultimately diffracted to a modest 3.7 Å resolution in the *P*_1_ space group consistent with 8 molecules (dimer of tetramers) in the asymmetric unit (ASU) (Table S1). This oligomerization is likely crystallization induced based on our in-solution studies (Fig. S1). We solved this structure by using molecular replacement coupled with model building and restrained refinement approaches (*R*_w_/*R*_f_ = 0.210/0.268) (Table S1), and an analysis of the crystal contacts suggested that the (His)_6_ tag may have impacted the crystal quality. To overcome this issue, *Vc*NFeoB was recloned into a vector encoding for N-terminal (His)_6_ tag fused to a cleavable SUMO moiety. Expression and purification followed the same procedures as previously described (*vide supra*), and the SUMO tag was cleaved prior to crystallization. Crystals of the SUMO-cleaved apo domain were generated and ultimately diffracted to 2.3 Å resolution in the *P*_121_ space group consistent with 2 molecules in the asymmetric unit (ASU) (Table S1). This oligomerization is likely crystallization induced based on our in-solution studies (Fig. S1). Using the 3.7 Å resolution model, we were able to solve the 2.3 Å resolution structure of the tagless protein (*R*_w_/*R*_f_ = 0.203/0.266) (Table S1).

The X-ray crystal structure of the apo *Vc*NFeoB NTPase domain reveals the presence of a typical NFeoB fold (Fig. 2). Distinctly present are the two common NFeoB subdomains: the globular G-protein subdomain that is responsible for binding and hydrolyzing nucleotides (31) (Fig. 2, blue) and the hammer-shaped guanine-dissociation inhibitor (GDI) subdomain that regulates nucleotide dissociation (32) and connects directly to the FeoB transmembrane region (Fig. 2, red). Within the G-protein domain, two switch regions (known as Switch I and Switch II) that regulate nucleotide hydrolysis and communicate nucleotide status to the GDI domain (31, 32) are present, albeit Switch I is mostly disordered, while Switch II is mostly ordered in the apo form. Surprisingly, comparison of the *Vc*NFeoB NTPase domain to other structurally characterized NFeoB domains displays strong structural conservation in both subdomains (C_α_ RMSD < 1Å on average) even though the NFeoB is not a strict GTPase (Fig. S2). These observations indicate that gross structural changes in the NFeoB region do not account for the observed nucleotide promiscuity of *V. cholerae* FeoB *per se*.

**Fig. 2.**
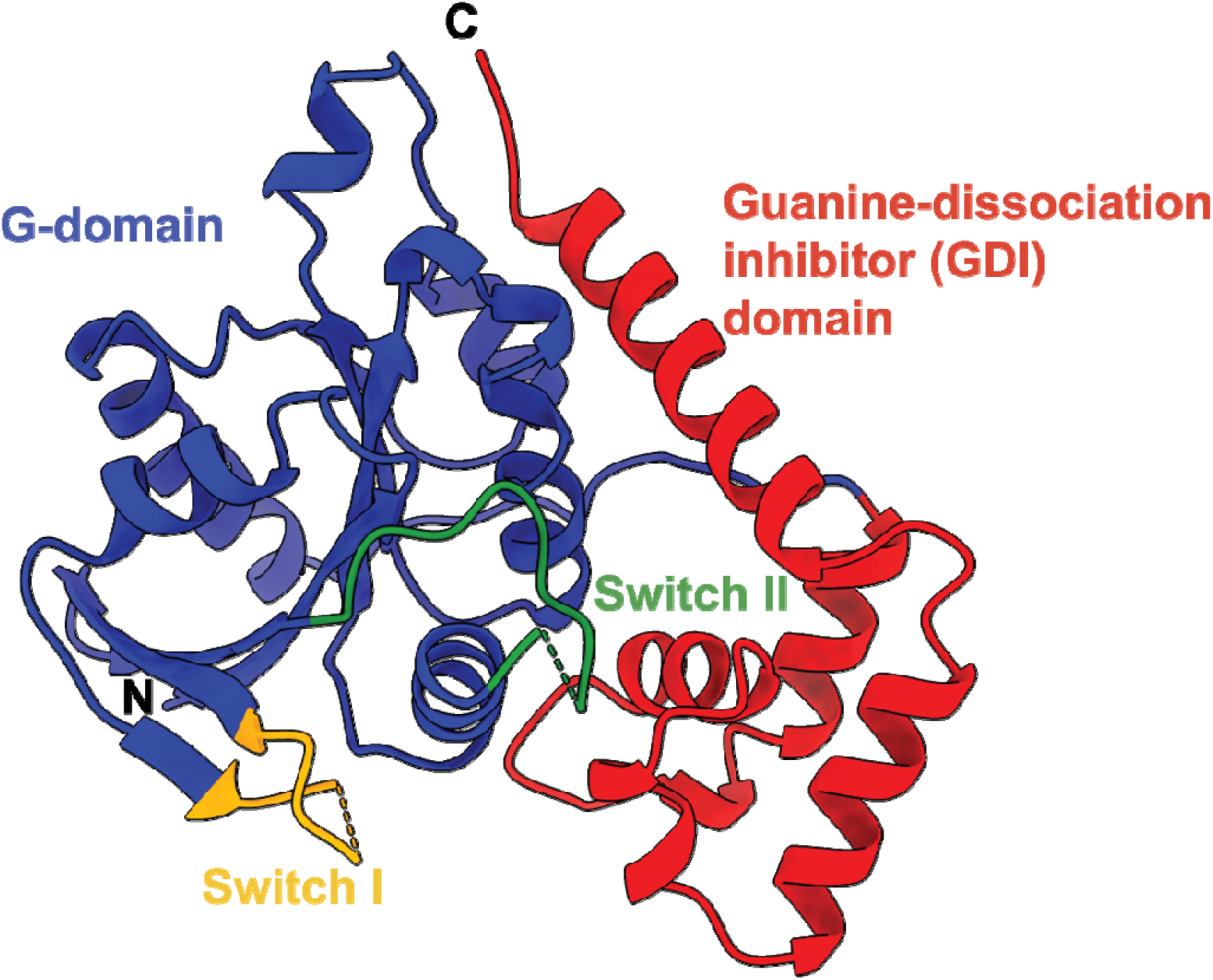
X-ray crystal structure of the SUMO-cleaved *Vibrio cholerae* NFeoB NTPase domain in the *apo* state (PDB ID 9BA6). The overall structure of the *Vc*NFeoB has a typical NFeoB fold and comprises two major domains: the guanine-dissociation inhibitor (GDI) domain (labeled in red) that regulates GDP release and connects to the transmembrane region, and the G-protein domain (labeled in blue) that binds and catalyzes nucleotide hydrolysis. Within the G-protein domain are two key switch regions (Switch I and Switch II, labeled yellow and green respectively) that regulate nucleotide hydrolysis and transmit information to the GDI domain. In the absence of nucleotide, Switch I is mostly disordered, while Switch II is mostly ordered. ‘N’ and ‘C’ represent the N- and C-termini in the structure, respectively.

### The GDP-bound Structure of the VcNFeoB NTPase Domain Reveals Important Nucleotide-Binding Residues

To determine whether structural properties within the nucleotide-binding pocket could contribute to nucleotide promiscuity, we sought to determine the structure of *Vc*NFeoB in the presence of various nucleotides. To do so, we co-crystallized apo *Vc*NFeoB (both SUMO-cleaved and (His)_6_ tagged) in the presence of hydrolyzed nucleotides (*e*.*g*., ADP and GDP) and the presence of non- or slowly-hydrolyzable triphosphate mimics (*e*.*g*., AMP-PNP, AMP-PCP, and GMP-PNP). Despite extensive fine screening and testing of multiple conditions, we were only able to generate datasets of GDP-bound *Vc*NFeoB(His)_6_ that diffracted modestly, but completely, to 4.2 Å resolution in the *C*_121_ space group consistent with 4 molecules in the asymmetric unit (ASU) (Table S1). This oligomerization is likely crystallization induced based on our in-solution studies (Fig. S1). Using molecular replacement with our 2.3 Å apo *Vc*NFeoB structure coupled with strongly restrained refinement approaches, we built a model that clearly displayed GDP in the nucleotide-binding pocket of all molecules in the ASU based on omit maps (Fig. S3). Using the X-ray structure of GDP-bound *E. coli* NFeoB as a guide (PDB ID 3I8X; 2.3 Å resolution), we were able to place GDP in all *Vc*NFeoB molecules in the ASU and to solve this structure (*R*_w_/*R*_f_ = 0.223/0.273) (Table S1).

The crystal structure of *Vc*NFeoB(His)_6_ bound to GDP reveals multiple amino acids that contribute to nucleotide binding (Fig. 3). In the absence of nucleotide, the binding pocket is fairly open, while binding of GDP elicits a contraction surrounding the nucleobase with the associated amino acids responsible for nucleotide recognition coming together (Fig. S4). In this structure, the Switch I loop that has been shown to be important for GTP hydrolysis (31) is mostly disordered, which may be attributed to the flexibility of this region when GDP is bound. As the nucleobase enters the binding pocket, a region of random coil from Asn119 to Asp122 tightens (Fig. S4), and both residues (conserved amongst NFeoBs) become within hydrogen-bonding distance (3.5 and 2.8 Å, respectively) of the guanine purine (Figs. 3,4a). Underneath the guanine purine is Lys120, also conserved in many NFeoBs, that appears to prop up the hydrolyzed nucleotide in the binding pocket; electrostatic contributions from this residue are not observed in our structure, and in some NFeoBs (like those in *S. thermophilus, K. pneumoniae, E. coli* and *Gallionella capsiferriformans*, and *Thermotoga maritima*) this residue corresponds to a non-polar amino acid, either a Met or an Ala(31, 33-36). Interestingly, Asn150 is positioned along a region of random coil (known as the G5 motif) above the guanine purine but turned towards, and tightly hydrogen bonded with Asp122 on the G4 motif (2.3 Å distance) (Figs. 3,4a). Because of the high variability within the G5 region, we sought to use bioinformatics to gain a better understanding of whether Asn150 might have an important role in nucleotide discrimination.

**Fig. 3.**
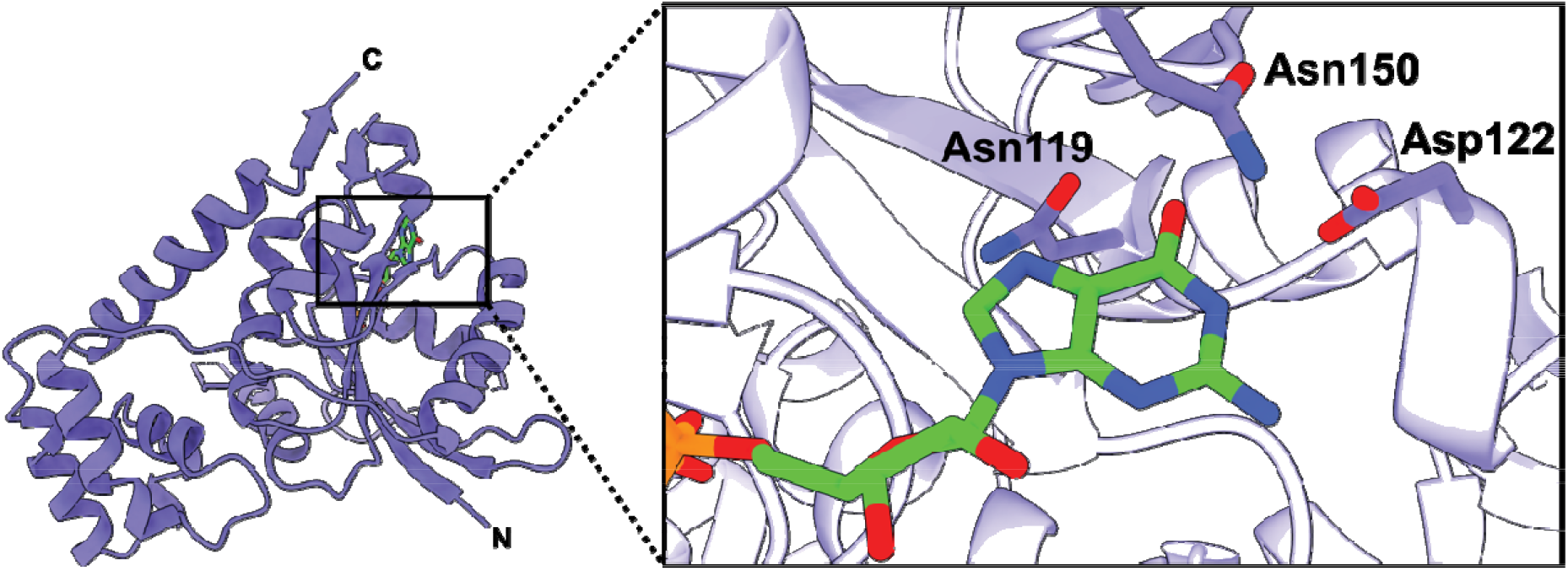
X-ray crystal structure of the *Vibrio cholerae* NFeoB NTPase domain in the GDP-bound state. The right panel represents a zoomed-in view of the nucleotide-binding pocket bound to GDP. Three residues make important contacts with the purine ring: Asn119, Asp122, and Asn150. Of these residues, only Asn150 is located on a variable loop region (G5) and lacks conservation. In the presence of GDP, the Switch I region is nearly fully disordered, while the Switch II region is only partially disordered. ‘N’ and ‘C’ represent the N- and C-termini in the structure, respectively.

**Fig. 4.**
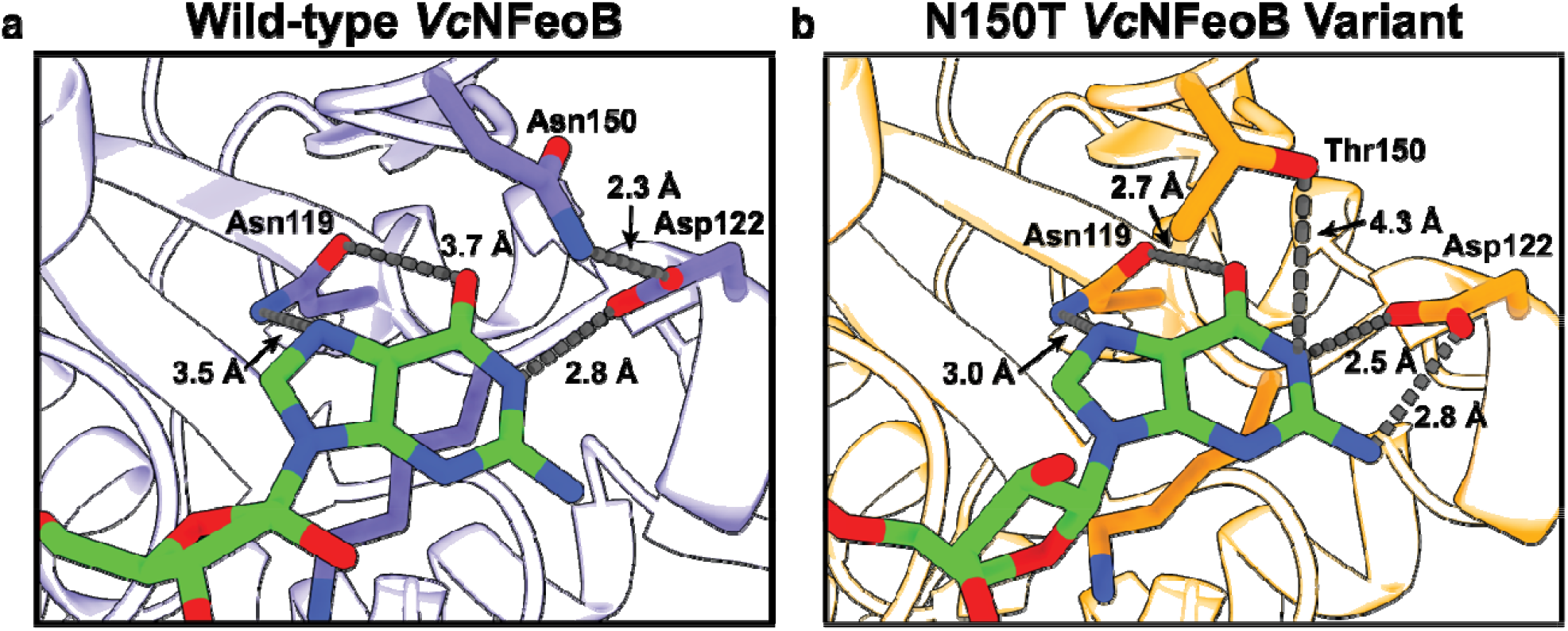
Comparisons of the nucleotide-binding pockets of the WT (**a**) and N150T (**b**) *Vc*NFeoB NTPas domain structures in their GDP-bound forms. Fewer hydrogen-bonding interactions are observed in the WT GDP-bound structure compared to the N150T variant of the NTPase domain. We hypothesize that th fewer hydrogen-bonding interactions in the WT structure allow for greater plasticity in NTP/NDP binding, unlike the strict GTPases that do not bind and hydrolyze other NTPs.

### Alterations in Hydrogen-Bonding Surrounding the Nucleotide-Binding Pocket Are Likely Linked To the Nucleotide Promiscuity of VcNFeoB

To gain a better understanding of residues that are either conserved or variable amongst NFeoB NTPases and GTPases, and to understand whether these sequence differences might contribute to functional differences, we used multiple sequence analysis (MSA) and phylogenetics. To do so, we limited our approach only to the sequences that have been previously tested *in vitro* to have either NTPase or strictly GTPase activities(28). Partial sequence alignments revealed that the position analogous to Asn150 within the G5 motif of the *Vc*NFeoB NTPase domain is highly variable for NFeoBs known to be NTPases, while this same position is an invariant Thr residue for NfeoBs known to be strict GTPases (Fig. 5A). The flanking regions of the G5 motif are also highly variable in NFeoB NTPases, while the same regions are highly conserved in NFeoB GTPases (Fig. 5A), suggesting that a degree of flexibility near the nucleobase may be important for nucleotide promiscuity. Intriguingly, phylogenetic analyses using the full-length sequences of *bona fide* FeoB NTPases and GTPases reveal a differential clustering among GTPase and NTPase proteins (Fig. 5B). While the number of FeoB proteins with verified nucleotide activity is low, this analysis could suggest that convergent evolution (and/or horizontal gene transfer) may have occurred amongst these organisms, similar to a previous hypothesis for the Feo system(13).

**Fig. 5.**
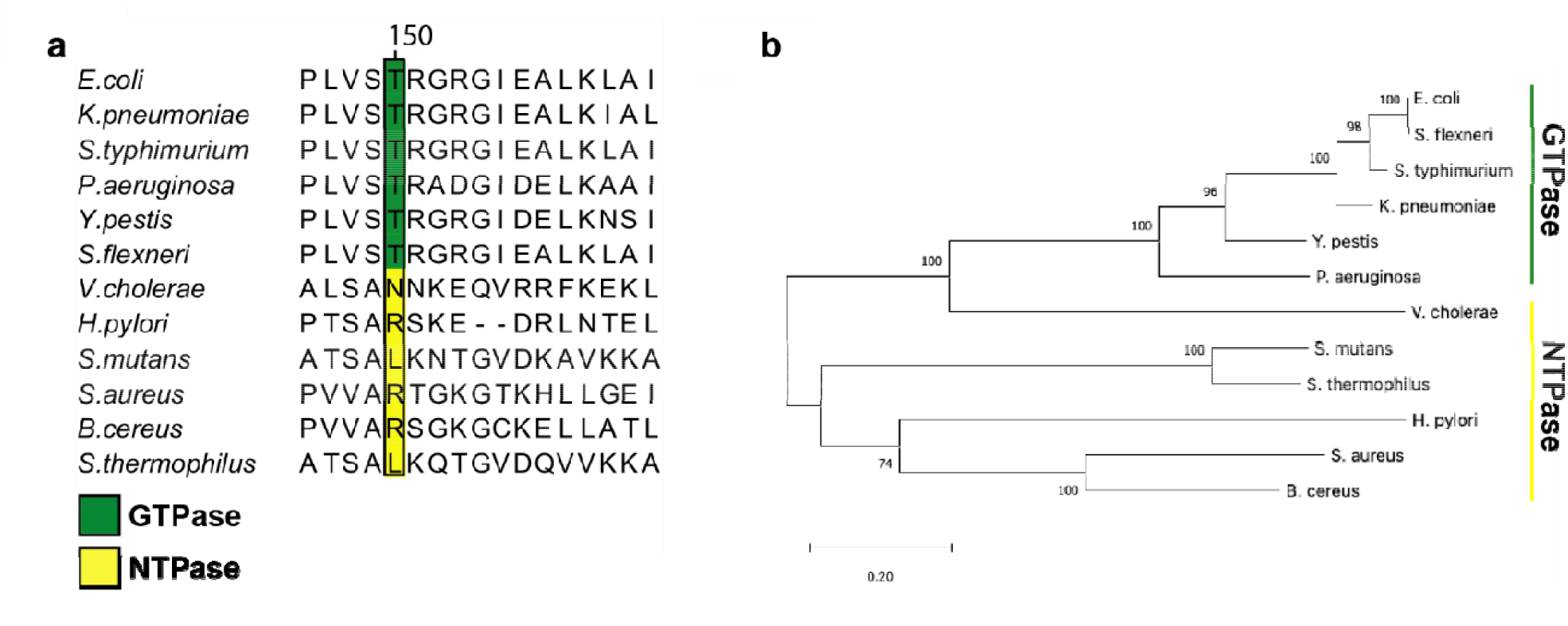
Partial multiple sequence alignment (MSA) and phylogenetic analysis of nucleotide-specific and nucleotide-promiscuous FeoBs. **a**. The partial MSA compares residues in the variable G5 region of experimentally determined GTPases (green) and experimentally determined NTPases (yellow). At position 150 in *V. cholerae* FeoB (NTPase) is an Asn residue, while the analogous position 150 in the strict GTPase FeoBs is a Thr residue. The numbering above the MSA is based on the *V. cholerae* FeoB sequence. **b**. Phylogenetic analysis of the FeoB sequences from the respective organisms in panel ‘**a**’ shows distinct clustering of the NTPase FeoBs from those of the GTPase FeoBs. The 0.20 scalebar indicates the amount of genetic change at a certain length. The numbers above the nodes represent the bootstrap values above 50% from 500 iterations.

To test whether alteration of Asn150 to a Thr residue would inform on structural changes that may take place within the nucleotide-binding pocket, we expressed, purified, and crystallized the N150T variant of *Vc*NFeoB(His)_6_ in the presence of GDP. The crystals diffracted to 2.9 Å resolution (Table S1), and initial analysis revealed one apo *Vc*NFeoB(His)_6_ NTPase domain and one GDP-bound *Vc*NFeoB(His)_6_ NTPase domain both within the asymmetric unit. This hetero-oligomerization is likely crystallization induced based on our in-solution studies (Fig. S1). Omit maps confirmed that substantial density was present and consistent with GDP in one, but not both, molecules within the asymmetric unit (Fig. S3), allowing us to visualize a direct comparison between the apo and the GDP-bound N150T variant at the same resolution (Fig. 6). Interestingly, the presence of Thr in position 150 affects the hydrogen bonding pattern surrounding the guanine nucleobase both directly and indirectly. First, the Thr hydroxyl moiety now makes a new hydrogen bond with position N1 along the purine ring (Fig. 4b). Second, removal of Asn150 releases Asp122 to tighten and to extend its hydrogen bonds with positions N1 and the H_2_N-C1 group (Fig. 4b). Third, Asn119 appears to pull the guanine further into the binding pocket, although uncertainty due to modest resolution of the GDP-bound WT *Vc*NFeoB NTPase domain structure prevents a definitive statement regarding Asn119 hydrogen bonding strength. However, by comparison to the WT protein, the N150T variant appears to have two key additional hydrogen bonds surrounding the guanine purine that could affect nucleotide stability and may afford the discrimination of GTP relative to other NTPs. Finally, like the WT *Vc*NFeoB NTPase domain, the N150T variant displays a mostly disordered Switch I region and a partially disordered Switch II region in both the GDP-bound and apo forms.

**Fig. 6.**
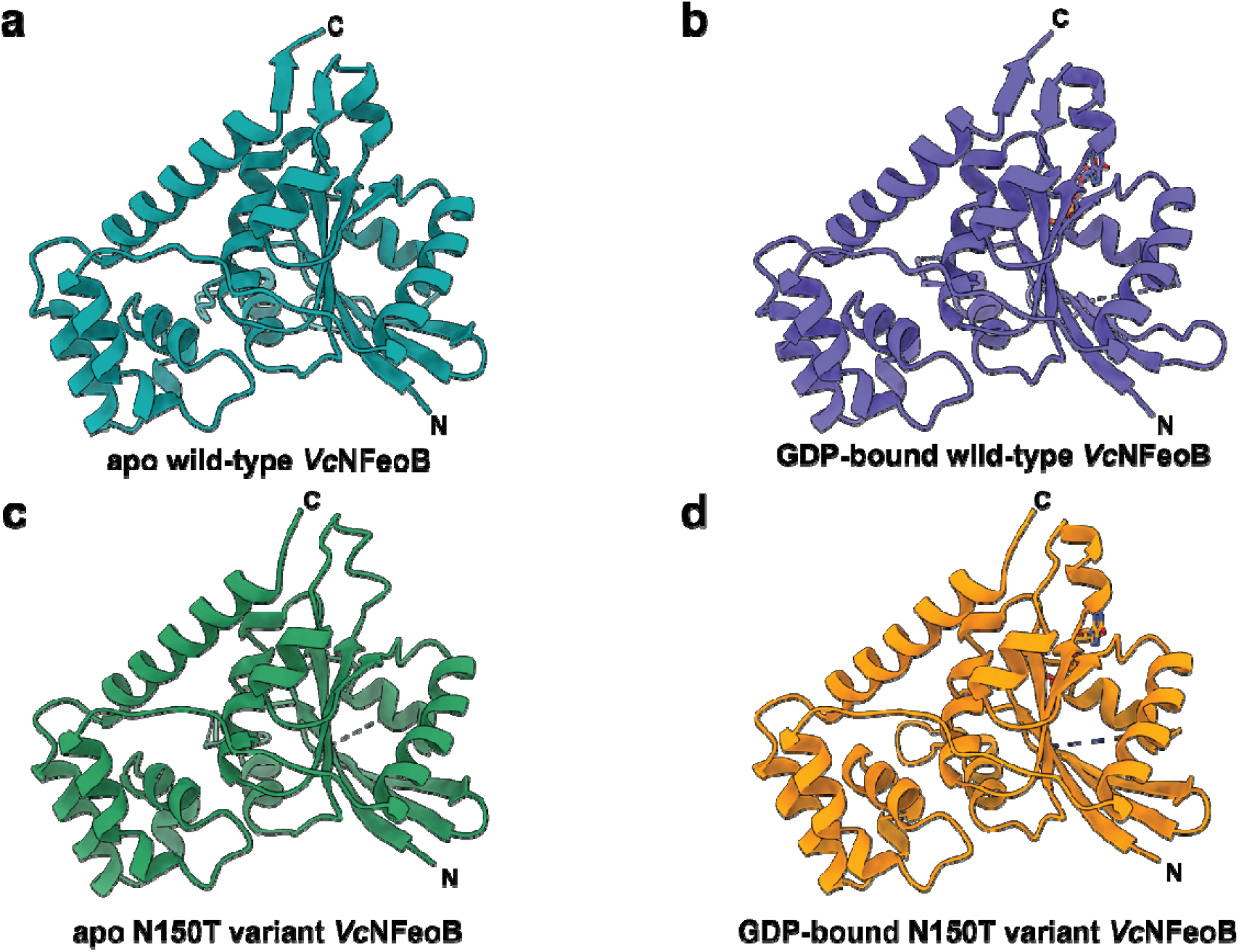
Comparisons of the WT and N150T *Vc*NFeoB NTPase domain structures in their apo and GDP-bound forms. **a**. Apo WT *Vc*NFeoB(His)_6_ NTPase domain (PDB ID 8VWL). **b**. GDP-bound WT *Vc*NFeoB(His)_6_ NTPase domain (PDB ID 8VWN). **c**. Apo N150T *Vc*NFeoB(His)_6_ NTPase domain (PDB ID 9BA7). **d**. GDP-bound N150T *Vc*NFeoB(His)_6_ NTPase domain (PDB ID 9BA7). In general, the structural similarities among the WT and variant *Vc*NFeoB(His)_6_ NTPase domains are very high: 1.36 Å C_α_ RMSD (apo WT and apo N150T) and 0.88 Å C_α_ RMSD (GDP-bound WT and GDP-bound N150T). ‘N’ and ‘C’ represent the N- and C-termini in the structure, respectively.

### Isothermal Titration Calorimetry Coupled with AlphaFold Modeling Reveal Key Differences in GTP/GDP and ATP/ADP Binding

To test the binding strength and binding stoichiometry of various nucleotides to the WT *Vc*NFeoB NTPase domain and its N150T variant, we used isothermal titration calorimetry (ITC) (Fig. 7). Despite the hydrogen-bonding differences in our X-ray crystal structures, the WT and N150T *Vc*NFeoB proteins displayed nearly identical binding strengths to GDP: WT K_*d*_ of 3.18 μM ± 0.23 μM; N150T K_*d*_ of 3.60 μM ± 0.36 μM (Fig. 7a,b) with a single binding site (N ≈ 1). By comparison to the binding of GMP-PNP (a non-hydrolyzable GTP analog), the WT and N150T *Vc*NFeoB constructs displayed slightly different binding strengths, consistent with our observed increase in hydrogen bonding in the N150T variant: WT K_*d*_ of 121.23 μM ± 34.13 μM; N150T K_*d*_ of 93.95 μM ± 15.63 μM (Fig. 7c,d) with a single binding site (N ≈ 1). GDP is known to have a stronger affinity to NFeoBs than GTP/GMP-PNP due to the presence and function of the GDI domain(32, 37), and we observed a similar trend as our structural work also reveals the presence of a GDI domain in *Vc*NFeoB. However, based on these data, the presence/absence of Asn150 alone does not dramatically change the binding strength of GTP/GDP to the *Vc*NFeoB NTPase domain. We then tested the ability of the *Vc*NFeoB NTPase domain to bind adenosine containing nucleotides. While we attempted multiple different concentrations and stoichiometries using both hydrolyzed (ADP) and non-hydrolyzable analogs (AMP-PNP), we did not observe any appreciable saturation that could be fitted to any logical binding isotherm for both the WT and N150T variant proteins (Fig. S5). However, titrations of ADP/AMP-PNP showed strong amounts of heat evolution (up to 12 μW per injection) even when corrected for dilutions of nucleotide in the absence of protein (Fig. S5). These observations suggest either a rapid kinetic reversibility (*i*.*e*., fast *k*_on_ and fast *k*_off_), and/or a potential conformational change that may be occurring as the domain samples ADP/AMP-PNP in the binding pocket.

**Fig. 7.**
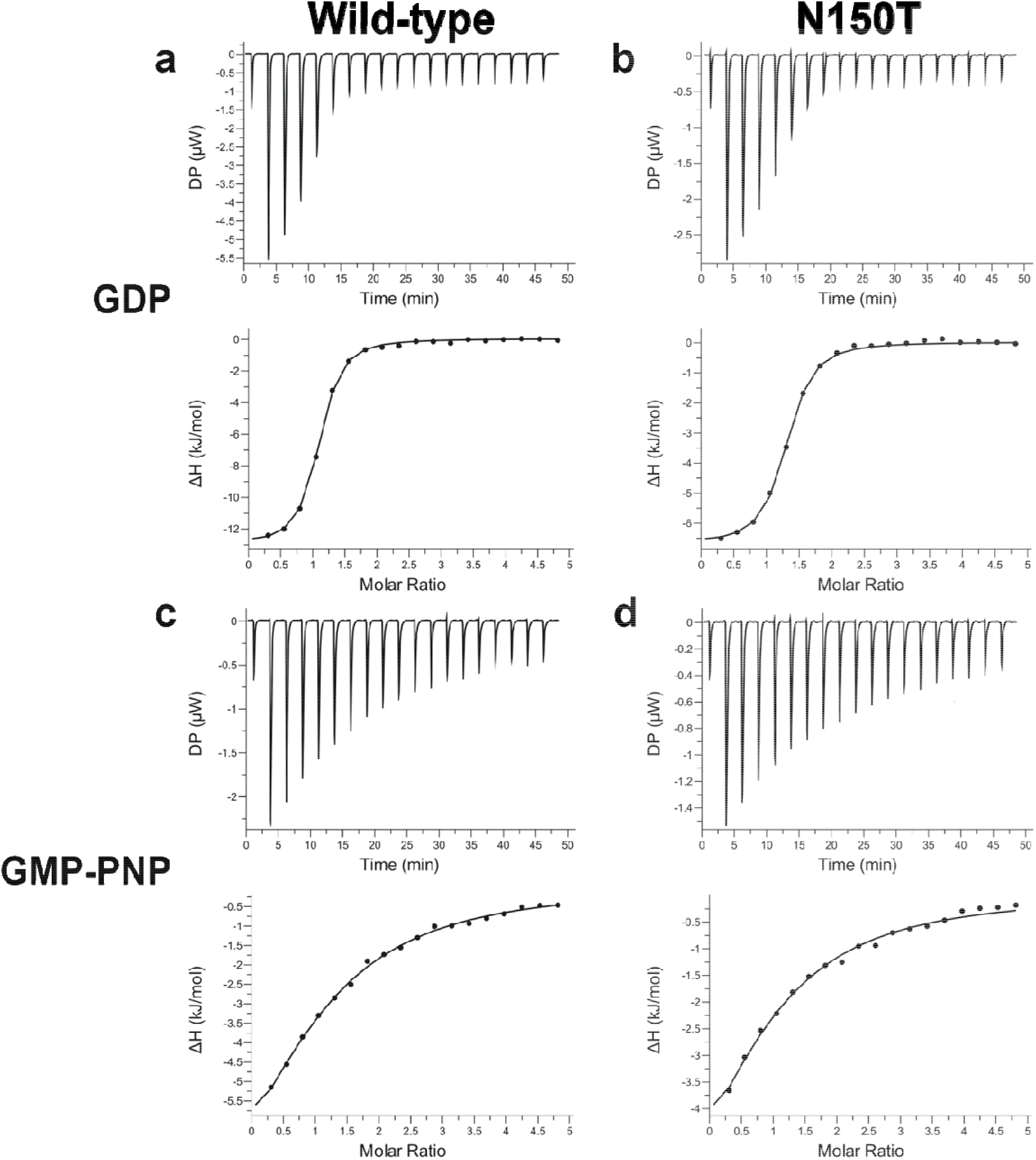
The *Vc*NFeoB NTPase domain binds GDP and GMP-PNP similarly to GTP-specific NFeoB domains. Representative ITC thermograms (top) and ΔH vs. molar ratio traces (bottom) of WT (left) and N150T (right) *Vc*NFeoB titrated with either GDP (**a, b**) or GMP-PNP (**c, d**). All datasets have been corrected for nucleotide dilution into buffer in the absence of protein. The strength of binding for GDP to both WT (3.18 μM ± 0.23 μM) and N150T *Vc*NFeoB (3.60 μM ± 0.36 μM) are nearly identical, with both having a single binding site (N ≈ 1). The strength of binding for GMP-PNP to WT *Vc*NFeoB (121.23 μM ± 34.13 μM) is slightly weaker than binding to N150T *Vc*NFeoB (93.95 μM ± 15.63 μM), but both reveal a single binding site (N ≈ 1). All values were determined in triplicate and represent the mean ± one standard deviation of the mean.

Finally, to gain a better understanding of what may be occurring within the binding pocket, we used the AlphaFold3 server to predict the WT and N150T *Vc*NFeoB structures bound to ADP and ATP (Fig. S6). Interestingly, in the predicted WT structures in the presence of adenine-containing nucleotides, Asn150 makes a very weak interaction with the adenine purine while Asp122 and Ser148 (part of the G4 and G5 motifs, respectively) turn completely away from the binding pocket and make tight hydrogen-bonds (≈ 2.7 Å) with one another (Fig. S6a). A similar result was observed in the N150T variant with Thr150, making even weaker interactions with the adenine nucleotide (Fig. S6b). In both cases, strong hydrogen-bond interactions (≤ 2.8 Å) with Ser148 cause the NTPase domain to adopt an “Asp off” conformation, *i*.*e*., Asp122 makes no interactions with the adenine nucleobase. These results differ from the GDP-bound structures, as both adopt a “Asp on” conformation in which Asp122 makes strong interactions (≈ 2.4 Å – 2.8 Å) with the guanine nucleobase. When we compare the ADP/ATP-bound behavior to the apo structure of *Vc*NFeoB (Fig. S4), we note that the G4 and G5 regions would need to undergo structural rearrangements upon nucleotide binding, which could explain the strong heat evolution upon ADP and AMP-PNP titrations, but fewer hydrogen bonding interactions likely preclude stable or prolonged binding within the pocket, explaining the ITC results.

## DISCUSSION

Whether FeoB, the primary prokaryotic ferrous iron transporter, is nucleotide promiscuous or nucleotide specific has vacillated for some time. Before the structure of NFeoB was known, early studies of FeoB predicted that the protein might hydrolyze ATP due to its sequence similarity to other ATPases, and decreased FeoB-dependent iron uptake in *Helicobacter pylori* was observed when ATP synthesis was disrupted by proton uncouplers(38-40). However, subsequent studies of FeoB showed that the NFeoB domain from *E. coli* was GTP-specific(41), and structures of NFeoB from *Methanocaldococcus jannaschii* and *E. coli* revealed the presence of a G-protein like domain(34, 42), strongly implying that NFeoB bound and hydrolyzed only guanine nucleotides. This presumption continued for nearly two additional decades, as additional NFeoB structures were determined and FeoB was further explored in an almost GTP-exclusive manner(12, 14, 18). However, despite this assumption, Shin *et al* reexamined NTPase activity in the context of *V. cholerae* FeoB and found this protein to be nucleotide promiscuous both *in vitro* and *in vivo(24)*. These observations were further expanded to show that several bacterial FeoBs could be differentially classified as GTP-specific while others could be classified as nucleotide promiscuous(24, 25). However, the structural basis for this functional divergence of FeoB was not known.

In this work, we provided the first X-ray crystal structure of *V. cholerae* NFeoB, a notably promiscuous NTPase, in its WT and variant forms, both in the presence and absence of nucleotides. While the general NFeoB G-protein like fold is conserved in our structures, the GDP-bound WT *Vc*NFeoB(His)_6_ structure revealed interactions between Asn150 (G5 motif) and Asp122 (G4 motif) that caused Asp122 to decrease the number of interactions it makes with the guanine nucleobase. As Asn150 is highly variable among NFeoB NTPases but conserved as a Thr among NFeoB GTPases, we wondered whether alteration of this residue alone could change nucleotide binding strength and/or specificity. Our structure of N150T *Vc*NFeoB(His)_6_ revealed increased H-bonding to the nucleobase due to the presence of Thr150 in the G5 motif, but this modification alone did not dramatically alter the binding affinity between NFeoB and GDP and only modestly increased the binding affinity between NFeoB and GMP-PNP. Instead, we hypothesized that the observed altered interactions might contribute more to nucleotide promiscuity.

While we were unable to determine experimental structures of adenine-based nucleotides bound to *Vc*NFeoB, ITC data and AlphaFold modeling provided insight into nucleotide promiscuity when taken in context of our other experimentally-determined structures. For example, ITC analyses of ADP- or AMP-PNP titrated into *Vc*NFeoB revealed an isotherm that failed to saturate but displayed strong heat evolution even after correction for appropriate dilutions, distinct from the response of this domain in the presence of GDP and GMP-PNP. This unusual behavior could be interpreted to mean weak binding but also that conformational changes may accompany the interactions of ADP/AMP-PNP within the nucleotide binding pocket, perhaps explaining why we failed to produce crystals of adenosine nucleotides bound to *Vc*NFeoB despite exhaustive trials. Structural modeling using AlphaFold supports this notion, as predicted structures of both ADP- and ATP-bound *Vc*NFeoB revealed movement of the G4 Asp122 residue and G5 Ser148 residue to lock the two amino acids into a strong H-bond preventing Asp122 from interacting with the nucleobase (*i*.*e*., “Asp off”). This conformational change opens up the binding pocket and results in only weak interactions as ADP/ATP enters but then exits the domain, likely rapidly. However, the rate of this conformational change must be slower than the rate of ATP hydrolysis, as we note that *Vc*NFeoB still hydrolyzes ATP robustly under these conditions(24, 25). In fact, all three residues (Asp122, Ser148, and Asn150) combined displayed an important role for ATP hydrolysis(24), suggesting the G4 and G5 motif are working in concert to contribute to nucleotide specificity.

Interestingly, structural analyses of other bacterial NTPases reveal a similar amino acid pattern that may contribute to a more general nucleotide promiscuity (Fig. S7), which could be leveraged for a functional advantage by select organisms. For example, among these NTPases in the G5 motif position analogous to Asn150 in *Vc*NFeoB are a diverse set of amino acids that only interact with the nucleotide base weakly at best(43), but this weak interaction may be important for plasticity within the binding pocket. In contrast, the positions analogous to Asp122 and Ser148 in *Vc*NFeoB are conserved in these other bacterial NTPases. Conservation of Asp in this region is unsurprising, as the G4 NxxD motif is required for GTP hydrolysis(44), but Ser is not; it may be possible that the Ser residue is needed to stabilize the “Asp off” conformation for NTPases to facilitate promiscuity in general. Based on these observations, we propose that a combination of these three amino acids in G4 and G5 provide conformational flexibility to allow the utilization of both GTP and ATP for protein function. In the case of FeoB, it is possible that this nucleotide promiscuity may provide an advantage to the several bacterial pathogens that are NTP agnostic. For these organisms, we propose that FeoB could leverage ATP hydrolysis when GTP levels in the cell are low, perhaps as a virulence factor or another adaptive mechanism to ensure Fe^2+^ uptake across a wide array of conditions, given iron’s essential nature to bacterial function. However, under homeostatic conditions, NTP promiscuous FeoBs likely rely on GTP and are likely still regulated by GDP based on the stable interactions observed in this study. Moreover, given the highly reactive nature of its translocated substrate, regulation of FeoB-mediated Fe^2+^ uptake by the status of GDP/GTP is presumably an important protective mechanism to prevent iron overload within the cell. It is possible that this intriguing mechanism of FeoB function could be leveraged as a means to combat bacterial virulence in the future.

## EXPERIMENTAL PROCEDURES

### Cloning of NFeoB Constructs

The *Vc*NFeoB(His)_6_ WT and N150T variant constructs were cloned into the pET-21a(+) plasmid as described previously (24) based on the sequence of WT *Vc*FeoB (Uniprot ID C3LP27). To create the N-terminal (His)_6_-SUMO-*Vc*NFeoB fusion, the gene encoding for *Vc*NFeoB was subcloned, and a synthetically added sequence for the Small Ubiquitin-like Modifier (SUMO) protein (Uniprot ID Q12306) was commercially appended (GenScript). The entire sequence was then subcloned into the pET-45b(+) plasmid between the *Pml*I and *Pac*I restriction sites, which allows for the translation of the N-terminal (His)_6_-SUMO-*Vc*NFeoB fusion when read in frame.

### Expression of NFeoB Constructs

The *Vc*NFeoB(His)_6_ WT and N150T variant plasmids were separately transformed into BL21(DE3) electrocompetent cells via electroporation, plated on Luria-Bertani (LB) plates supplemented with ampicillin (100 μg/mL), and incubated at 30 ºC overnight. The next day, starter flasks containing 100 mL LB broth and ampicillin (100 μg/mL) were inoculated with a single colony each (WT and N150T) and allowed to grow overnight at 30 ºC with 200 RPM shaking. The next day, 25 mL of the overnight cultures were inoculated into 1 L flasks charged with 1 L LB broth and ampicillin (100 μg/mL, final), and these cells were grown at 37 ºC with shaking of 200 RPM. When OD_600_ reached 0.4-0.8, the cells in the flasks were then cold shocked at 4 ºC for 2 h before induction with isopropyl β-D-1-thiogalactopyranoside (IPTG) to a final concentration of 1 mM. Cells were then grown for *ca*. 20 h overnight and harvested the next day by spinning at 5000 x*g*, resuspended in resuspension buffer (50 mM Tris pH 8.0, 300 mM NaCl, and 10 % (v/v) glycerol) before being flash frozen on N_2*(l)*_ and stored at - 80 ºC.

All purification steps were conducted at 4 ºC unless otherwise stated. Frozen cells were thawed and diluted to 100 mL with resuspension buffer, and 1 mM (final) phenylmethylsulfonyl fluoride (PMSF) was added prior to sonication at 80 % amplitude, 30 s on pulse, 30 s rest pulse, 12 min total. Lysed cells were clarified by spinning at 163,000 x*g* for 1 h. The supernatant was then applied to a 5 mL HisTrap HP column (Cytiva) that was pre-charged with Ni^2+^ and equilibrated with 5 column volumes (CVs) of wash buffer (50 mM Tris pH 8.0, 300 mM NaCl, 10 % (v/v) glycerol and 1 mM TCEP). After the sample was applied, the column was washed with 10 CVs of wash buffer, then with wash buffer containing 50 mM imidazole, and eluted with wash buffer containing 150 mM imidazole. Eluted protein fractions were then pooled, and buffer exchanged into ion exchange wash buffer (50 mM Tris pH 8, 10 % (v/v) glycerol) using a 50 mL HiPrep 26/10 desalting column to remove all salt content before anion exchange chromatography. After desalting, the protein was then applied to a 5 mL HiTrap Q HP anion exchange column (Cytiva) that was then washed extensively with 10 CVs of the ion exchange buffer. The protein was purified via a linear elution gradient from 0 M to 1 M NaCl. Eluted protein fractions were pooled, concentrated via a 10 kDa molecular weight cutoff (MWCO) filter, and injected onto a 120 mL Superdex 75 preparative grade gel filtration column (Cytiva) after equilibration with 1.5 CVs of size exclusion buffer (25 mM Tris pH 8.0, 100 mM NaCl, 5 % (v/v) glycerol and 1 mM TCEP). Fractions corresponding to pure, monomeric *Vc*NFeoB(His)_6_ were pooled and concentrated via a 10 kDa MWCO filter to *ca*. 12 mg/mL, aliquoted, flash frozen on N_2*(l)*_ and stored at −80 ºC. An identical procedure was followed for the *Vc*NFeoB(His)_6_ N150T variant.

The cellular transformation, expression, cellular harvesting, cellular lysis, and initial Ni^2+^-based purification of the (His)_6_-SUMO-*Vc*NFeoB mirrored that of *Vc*NFeoB(His)_6_. After fractions were eluted from the 5 mL HisTrap HP column, the (His)_6_-SUMO-*Vc*NFeoB protein was buffer exchanged into SUMO cleavage buffer (50 mM Tris pH 8.0, 300 mM NaCl, 10 % (v/v) glycerol, and 10 mM β-mercaptoethanol (BME)) and concentrated via a 10 kDa MWCO filter. House-made SUMO protease Ulp1 was then added at a 1:100 (mg/mg) ratio and allowed to cleave overnight with gentle rocking at 4 ºC. The next day, the solution was applied again to a 5 mL HisTrap HP column to separate the now cleaved *Vc*NFeoB from any uncleaved protein. Eluted fractions containing cleaved *Vc*NFeoB were concentrated via a 10 kDa MWCO filter injected onto a 120 mL preparative Superdex 75 column, eluted isocratically, pooled, and stored identically to *Vc*NFeoB(His)_6_ (*vide supra*).

### Crystallization of NFeoB Constructs

Apo *Vc*NFeoB(His)_6_ was initially thawed and diluted to 10 mg/mL with size exclusion buffer prior to crystallization trials. Several commercial sparse-matrix screens were used to test for crystallization using vapor diffusion in sitting drop format. After incubation at 25 ºC for *ca*. 4 months, crystals appeared in a condition containing 25 % (w/v) PEG 3350, 0.1 M bis-Tris pH 5.5, and 0.2 M MgCl_2_. The crystals were then looped, cryo-protected, and frozen in N_2*(l)*_. Unfortunately, fine screens failed to replicate crystallization for further optimization.

To crystallize GDP-bound *Vc*NFeoB(His)_6_, protein at 10 mg/mL was incubated with 3 mM GDP for 2 h at room temperature before sparse-matrix screens were used to test for crystallization using vapor diffusion in sitting drop format at 25 ºC. After 2 weeks, small, cubic-shaped crystals appeared in a condition containing 25 % (w/v) PEG 3350, 0.1 M bis-Tris pH 5.5, 0.2 M MgCl_2_, and 0.1 M LiCl. Crystals were then looped, cryo-protected, and frozen in N_2*(l)*_.

The preparation of GDP-bound *Vc*NFeoB(His)_6_ N150T was identical to that of the WT protein. Sparse-matrix screens were used to test for crystallization using vapor diffusion in sitting drop format at 25 ºC. Crystals initially appeared in conditions containing 24 % (w/v) PEG 3350, 0.1 M bis-Tris pH 5.5, 0.03 M MgCl_2_, and 0.2 M (NH_4_)_2_SO_4_. Crystals matured after two weeks and were then looped, cryo-protected, and frozen in N_2*(l)*_.

To crystallize SUMO-cleaved *Vc*NFeoB, the protein was initially thawed and incubated with 3 mM ADP prior to dilution to 10 mg/mL. Sparse-matrix screens were used to test for crystallization using vapor diffusion in sitting drop format at 20 ºC. After two weeks of incubation, clustered crystals appeared in a condition containing 30 % (w/v) PEG 2000, 0.1 M Tris pH 8.0. Single crystals were separated manually using crystallization tools after the clusters were transferred to a drop containing cryo-protectant. Separated single crystals were then looped and frozen in N_2*(l)*_.

### X-ray Diffraction, Data Reduction, and Structural Determination

Diffraction data were collected at the Advanced Photon Source (APS), Argonne National Laboratory on LS-CAT beamline 21-ID-D and at Brookhaven National Laboratory beamline 17-ID-2 (FMX). Data were automatically processed using Xia2 (45) and/or AutoProc(46). The initial phases of all datasets were determined by molecular replacement (MR) using Phenix Phaser (47) with an AlphaFold-generated model as an initial search input(48). After an initial MR solution was identified, further model building was accomplished using Phenix AutoBuild(47). The unambiguous presence of GDP in the nucleotide-binding site was confirmed by the generation of Polder maps in Phenix for GDP-bound *Vc*NFeoB(His)_6_ WT and N150T datasets. The initial placement of GDP was determined based on the structure of *E. coli* NFeoB bound to GDP (PDB ID 3I8X) that was then further refined. Iterative rounds of manual model building and refinement were accomplished in Coot (49) and Phenix Refine(47), respectively, until model convergence and the final placement of any visible solvent molecules. Ramachandran statistics and clash values were determined from the MolProbity program (50) within the Phenix software suite. The following structures have been deposited in the Protein Data Bank: WT apo *Vc*NFeoB(His)_6_ (PDB ID 8VWL); WT GDP-bound *Vc*NFeoB(His)_6_ (PDB ID 8VWN); N150T GDP-bound *Vc*NFeoB(His)_6_ (PDB ID 9BA7); SUMO-cleaved apo *Vc*NFeoB (PDB ID 9BA6). Data collection and refinement statistics are provided for all structures in SI Table 1.

### Isothermal Titration Calorimetry

Purified WT or N150T *Vc*NFeoB(His)_6_ was diluted to 0.1 mM (3.2 mg/mL) in SEC buffer for all isothermal titration calorimetry (ITC) experiments. Experiments were conducted using the MicroCal PEAQ-ITC Automated (Malvern Panalytical) to probe nucleotide binding. All titrations with nucleotide diphosphates (GDP and ADP) were performed in 25 mM Tris pH 8.0, 100 mM NaCl, 5% (v/v) glycerol and 1 mM TCEP, while experiments involving the triphosphate mimics (GMP-PNP and AMP-PNP) were conducted using the same buffer conditions except with added MgCl_2_ to 10 mM final concentration. The calorimetry cell was loaded with 200 μL of WT or N150T *Vc*NFeoB(His)_6_, and 40 μL of nucleotide (GDP, GMP-PNP, ADP, or AMP-PNP) at 2.5 mM concentration (GDP and GMP-PNP) or 5.0 mM concentration (ADP and AMP-PNP) was loaded into the injection syringe. Thermal equilibrium was reached at 25 ºC after an initial 60 s delay followed by 19X 2 μL serial injections into the cell with 150 s interval delays between injection points with high spinning. Data were analyzed using the Malvern MicroCal PEAQ ITC analysis tool and fitted to a binding isotherm that has a single site using the following equations:

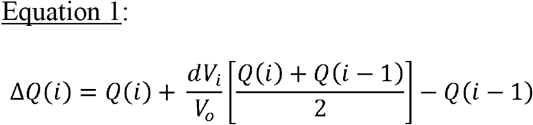

Where the heat released, ΔQ(i), from the ith injection is represented by ΔQ(i).

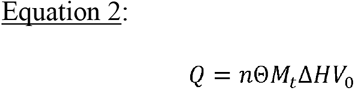

Where the total heat (Q) is related to the number of sites (n), the fractional occupation (Θ) the total free concentration of the macromolecule (*M*_t_), the molar heat of ligand binding (ΔH), and the volume determined relative to zero for the unbound species (V_0_).

### Bioinformatic Analyses

Based on previous studies in which nucleotide promiscuity of NFeoB from multiple organisms was initially uncovered(28), sequences were obtained from the Uniprot database of intact FeoBs. A multiple sequence alignment was constructed through the EMBL MUSCLE program using default parameters(51). The resultant alignment was visualized via Jalview (52-54) and was then entered into the MEGAX software (55-57) for phylogenetic analysis using the maximum likelihood method and 500 bootstrap iterations with a minimum coverage of 95%. The final phylogenetic results were also visualized using MEGAX.

### Structural prediction using AlphaFold3

Predicted *Vc*NFeoB structures with adenosine nucleotides (ATP and ADP) were generated using the AlphaFold3 (58) server by submitting amino acids 1-261 from *Vc*FeoB (Uniprot ID C3LP27) with either ADP and Mg^2+^ or ATP and Mg^2+^ and utilizing the default parameters. In all cases, the lowest energy calculated structure is displayed as being representative, but the resulting five calculated structures for each prediction reveal very similar results. The calculated structures were both visualized and analyzed using ChimeraX(59).

## Acknowledgements

This work was primarily supported by NSF CAREER grant CHE1844624 (A. T. S.), and in part by NIH-NIGMS grant R35 GM133497 (A.T.S), NIH-NIGMS grant T32 GM066706 (A. T. S. and M.L.), and by NIH-NIAID grants R01 AI091957 and R37 AI016935 (S.M.P.). This research used resources of the Advanced Photon Source, a U.S. Department of Energy (DOE) Office of Science User Facility operated for the DOE Office of Science by Argonne National Laboratory under Contract No. DE-AC02-06CH11357. Use of the LS-CAT Sector 21 was supported by the Michigan Economic Development Corporation and the Michigan Technology Tri-Corridor (Grant 085P1000817). NSLS2 is a U.S.DOE Office of Science User Facility operated under Contract No. DE-SC0012704. This publication resulted from the data collected using the beamtime obtained through NECAT BAG proposal # 311950. Sequence searches utilized both database and analysis functions of the Universal Protein Resource (UniProt) Knowledgebase and Reference Clusters (http://www.uniprot.org) and the National Center for Biotechnology Information (http://www.ncbi.nlm.nih.gov/).

## Supporting Information

Supporting information is available online, including:

Data collection and refinement statistics for apo WT*Vc*NFeoB(His)_6_ (*P*_1_ space group), GDP-bound WT *Vc*NFeoB(His)_6_, cleaved WT*Vc*NFeoB (*P*_121_ space group), and apo and GDP-bound N150T *Vc*NFeoB(His)_6_ (Table S1)

Size-exclusion chromatograms and 15 % SDS-PAGE analyses of purified *Vc*NFeoB constructs (Fig. S1)

Superpositioning of NFeoB domains (Fig. S2)

Quality of the electron density and omit maps surrounding the GDP molecule in WT *Vc*NFeoB(His)_6_ and N150T *Vc*NFeoB(His)_6_ (Fig. S3)

Comparison of the GTP-binding pocket of WT *Vc*NFeoB(His)_6_ in the apo state and the GDP-bound state (Fig. S4)

ITC data of adenine-containing nucleotides titrated into *Vc*NFeoB (Fig. S5)

AlphaFold3 predictions of *Vc*NFeoB bound to adenine nucleotides (Fig. S6

Altered hydrogen bonding interactions among various bacterial NTPases (Fig. S7)

## Data Availability Statement

All data are contained within the manuscript, either in the main body or in the Supplemental data submitted with the manuscript, and/or deposited in repositories such as the Protein Data Bank (PDB).

## Abbreviations

The abbreviations used are

GDP: guanosine diphosphate
GMP-PNP: 5’guanylyl-imidodiphosphate
GTP: guanosine triphosphate
ADP: adenosine diphosphate
AMP-PNP: 5’adenylyl-imidodiphosphate
ATP: adenosine triphosphate
IMAC: immobilized metal affinity chromatography
AIEX: anion exchange chromatography
IPTG: isopropyl β-D-l-thiogalactopyranoside
NFeoB: soluble N-terminal GTP-binding domain of FeoB
SUMO: small ubiquitin-like modifier
Ulp1: Ubl-specific protease
PMSF: phenylmethylsulfonyl fluoride
SDS-PAGE: sodium dodecyl sulfate polyacrylamide gel electrophoresis
SEC: size-exclusion chromatography
R.M.S.D.: root-mean-square deviation
Tris: tris(hydroxymethyl)aminomethane
TCEP: tris(2-carboxyethyl)phosphine
ITC: isothermal titration calorimetry;

