## Supporting Information for "Structural Determinants of *Vibrio cholerae* FeoB Nucleotide Promiscuity"

|  | Apo WT*Vc*NFeoB(His)_6_ in *P_1_* space group (PDB ID 8VWL) | GDP-bound WT*Vc*NFeoB(His)_6_ (PDB ID 8VWN) | Apo WT*Vc*NFeoB in *P_121_* space group (PDB ID 9BA6) | Apo and GDP-bound N150T *Vc*NFeoB(His)_6_ (PDB ID 9BA7) |
| --- | --- | --- | --- | --- |
| **Data Collection** |  |  |  |  |
| Beamline | APS 21-ID-D | APS 21-ID-D | NSLS-II 17-ID-2 | NSLS-II 17-ID-2 |
| Wavelength (Å) | 0.97918 | 1.12706 | 0.97934 | 0.97934 |
| Space group | *P_1_* | *C_121_* | *P_121_* | *P_21212_* |
| Cell Dimensions |  |  |  |  |
| *a*, *b*, *c* (Å) | 48.35, 84.12, 158.03 | 194.27, 155.31, 50.92 | 53.96, 73.44, 63.03 | 202.46, 50.09, 62.25 |
| α, β, γ (˚) | 76.17, 83.91, 74.51 | 90.00, 95.88, 90.00 | 90.00, 94.25, 90.00 | 90.00, 90.00, 90.00 |
| Resolution (Å) | 77.59-3.67 | 60.53-4.25 | 43.41-2.38 | 33.94-2.88 |
| *R*_merge_ | 0.122 (1.185) | 0.184 (0.912) | 0.130 (0.951) | 0.155 (1.907) |
| *CC_1/2_* | 0.998 (0.792) | 0.983 (0.356) | 0.992 (0.521) | 0.997 (0.330) |
| *I/σ(I)* | 7.0 (0.8) | 6.9 (0.9) | 11.4 (1.6) | 8.8 (0.9) |
| Completeness (%) | 95.0 (88.9) | 98.8 (87.8) | 99.9 (99.0) | 99.4 (100) |
| **Refinement** |  |  |  |  |
| Resolution (Å) | 77.57-3.67 (3.80-3.67) | 50.65-4.25 (4.41-4.25) | 39.46-2.38 (2.47-2.38) | 33.94-2.88 (2.98-2.88) |
| No. reflections | 24,186 | 10,396 | 19,694 | 14,940 |
| *R*_work_ | 0.210 | 0.223 | 0.203 | 0.214 |
| *R*_free_ | 0.268 | 0.273 | 0.266 | 0.268 |
| No. atoms/molecules |  |  |  |  |
| Protein | 15,757 | 7,888 | 4,101 | 3,962 |
| Mg^2+^ | 6 | 3 | 0 | 0 |
| GDP | 0 | 4 | 0 | 1 |
| Waters | 0 | 1 | 92 | 14 |
| Glycerol | 0 | 0 | 7 | 0 |
| Cl^-^ | 0 | 0 | 2 | 5 |
| SO_4_^2-^ | 0 | 0 | 0 | 6 |
| Average B-factors (Å^2^) | 147.0 | 150.0 | 48.0 | 85.0 |
| R.m.s. deviations |  |  |  |  |
| Bond lengths (Å) | 0.003 | 0.003 | 0.009 | 0.009 |
| Bond angles (˚) | 0.62 | 0.76 | 1.70 | 1.16 |
| Ramachandran plot |  |  |  |  |
| Favored | 95.2 % | 95.7 % | 97.6 % | 96.7 % |
| Allowed | 4.6 % | 4.1 % | 2.5 % | 3.3 % |
| Outlier | 0.2 % | 0.2 % | 0 % | 0 % |

**Table S1.** Data collection and refinement statistics for apo WT*Vc*NFeoB(His)_6_ (*P*_1_ space group), GDP-bound WT *Vc*NFeoB(His)_6_, cleaved WT*Vc*NFeoB (*P*_121_ space group), and apo and GDP-bound N150T *Vc*NFeoB(His)_6_. Parentheses indicate the highest resolution shells.

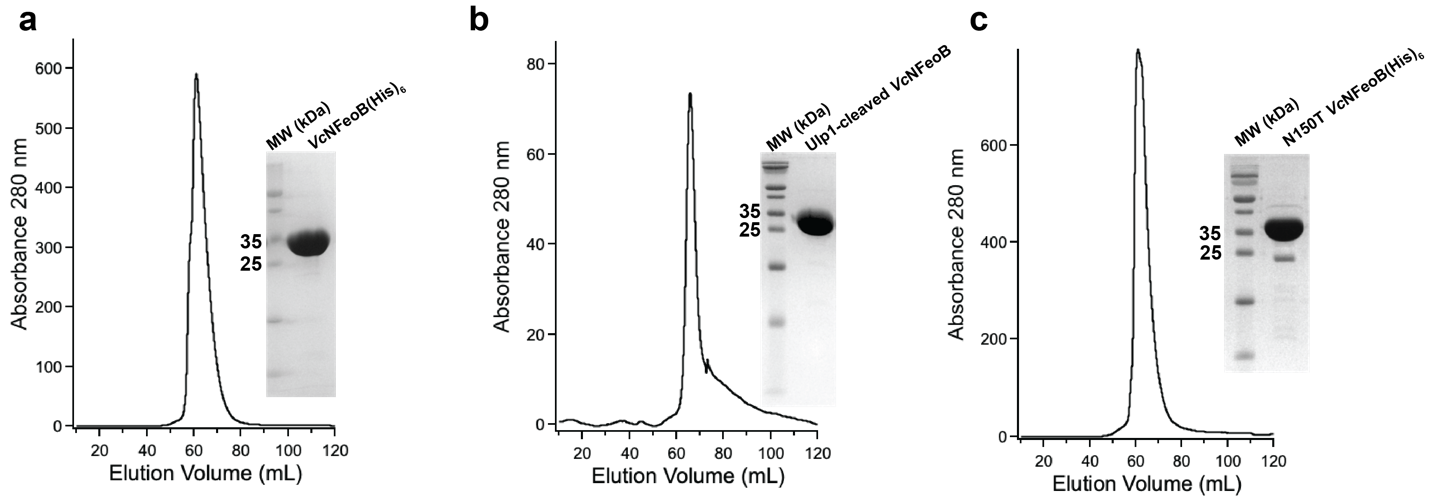

**Fig. S1**. Size-exclusion chromatograms and 15% SDS-PAGE analyses of purified *Vc*NFeoB constructs used in this work: **a**. WT *Vc*NFeoB(His)_6_; **b**. Ulp1-cleaved WT *Vc*NFeoB; **c**. N150T *Vc*NFeoB(His)_6_.

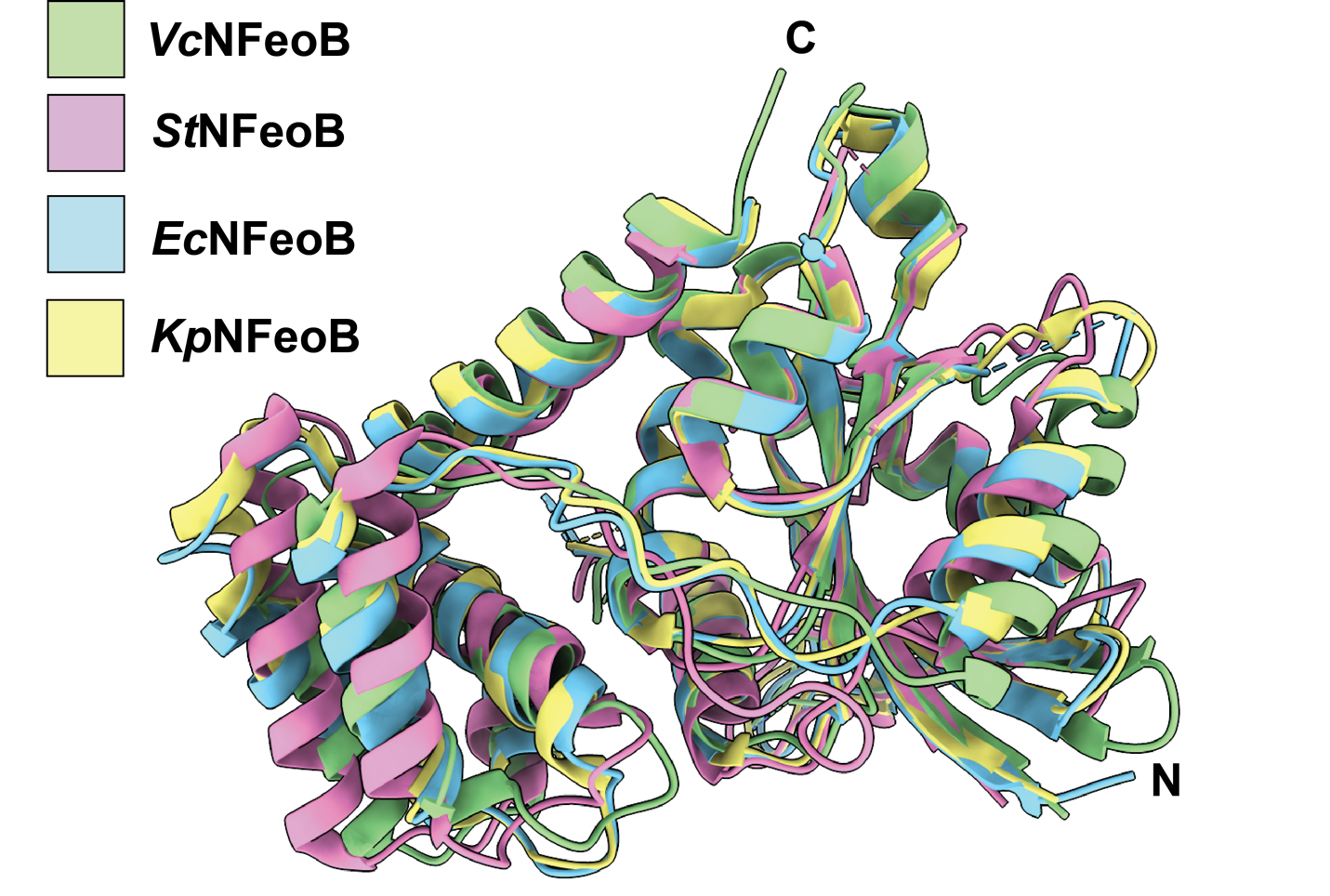

**Fig. S2.** Superpositioning of NFeoB domains in the apo state reveals structural conservation even though some domains are GTP specific while others are NTP promiscuous. The apo structure of *Vibrio cholerae* NFeoB (PDB ID 9BA6, green) superposed onto the apo NFeoB structures from *Streptococcus thermophilus* (PDB ID 3B1Z, pink), *Escherichia coli* (PDB ID 2WIA, blue) and *Klebsiella pneumoniae* (PDB ID 3HYR, yellow). In general, the C_α_ RMSD of each structure relative to *Vc*NFeoB is *ca.* 1 Å. ‘N’ and ‘C’ represent the N- and C-termini in the structure, respectively.

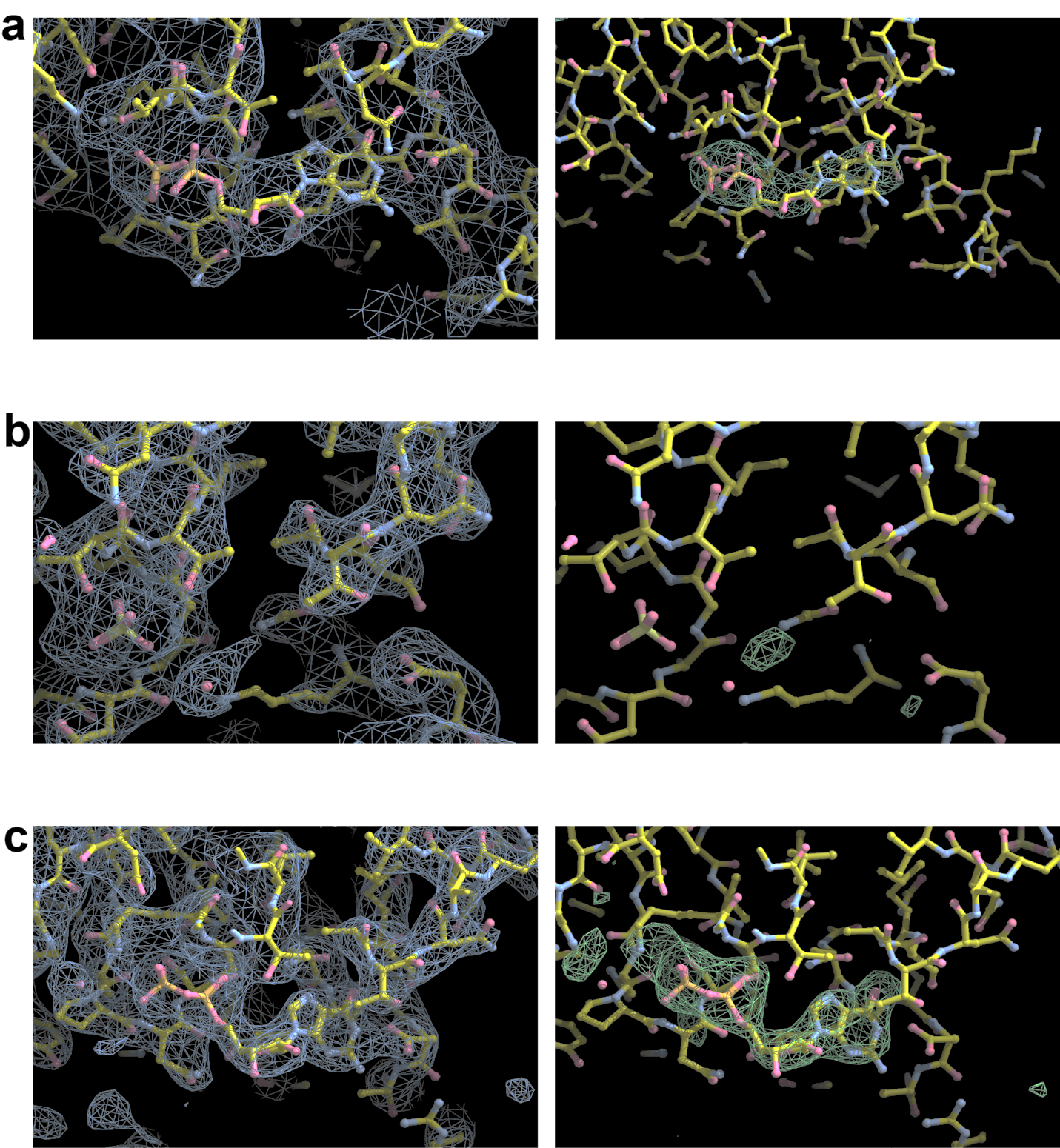

**Fig. S3**. Quality of the electron density and omit maps surrounding the GDP molecules in WT *Vc*NFeoB(His)_6_ and N150T *Vc*NFeoB(His)_6_. **a**. 2F_o_-F_c_ map of GDP-bound WT *Vc*NFeoB(His)_6_ on the left (contoured to 1σ), and the Polder omit map of GDP-binding pocket in the absence of GDP (contoured to 3 σ). **b**. 2F_o_-F_c_ map of the *Vc*NFeoB(His)_6_ N150T protomer that does not contain GDP, and the Polder omit map of the *Vc*NFeoB(His)_6_ N150T protomer that does not contain GDP in the GDP-binding pocket (contoured to 3 σ). **c.** 2F_o_-F_c_ map of the *Vc*NFeoB(His)_6_ N150T protomer that does contain GDP, and the Polder omit map of the *Vc*NFeoB(His)_6_ N150T protomer that does contain GDP in the GDP-binding pocket with GDP omitted (contoured to 3 σ).

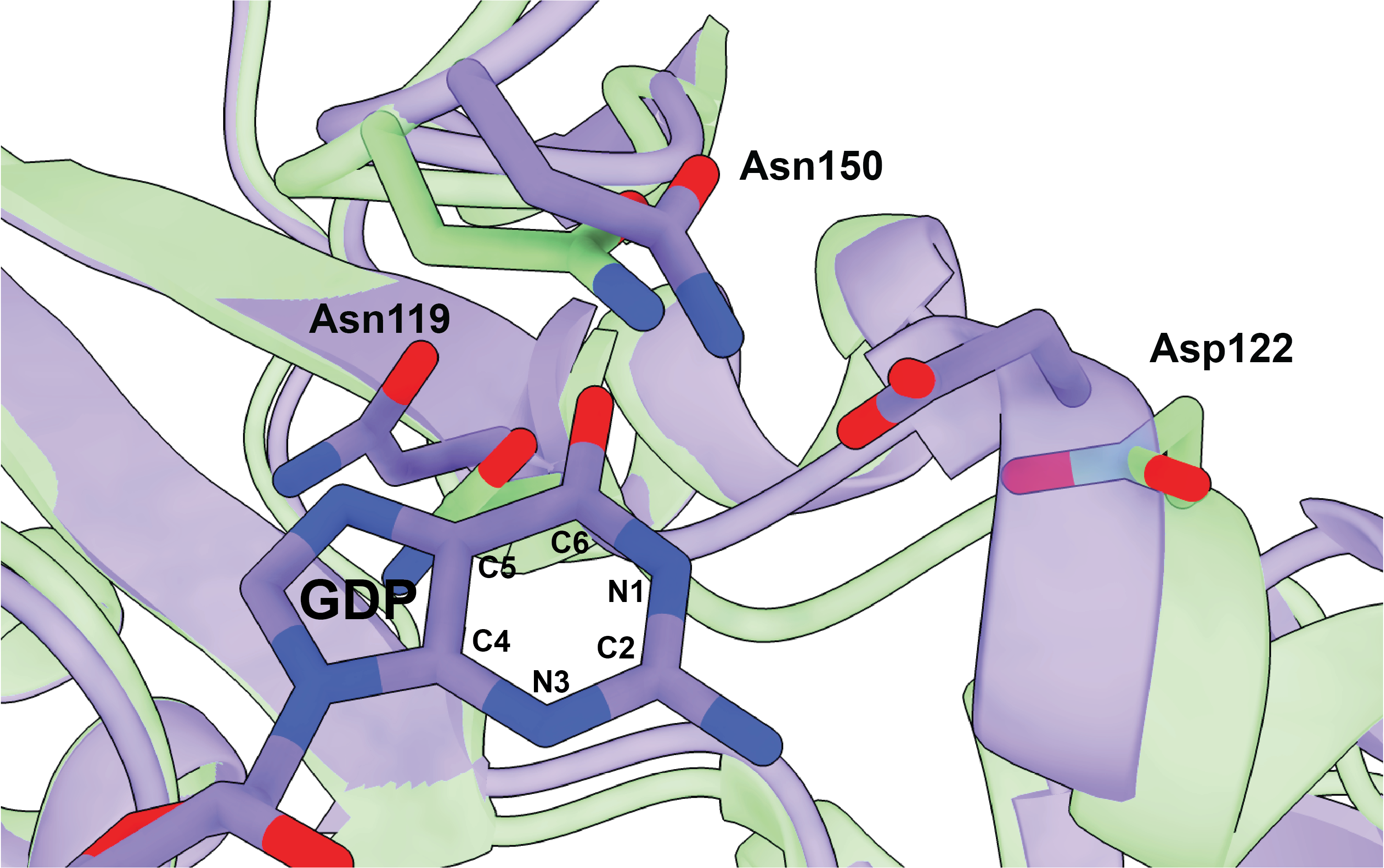

**Fig. S4.** Comparison of the GTP-binding pocket of WT *Vc*NFeoB in the apo state (green) and the GDP-bound state (purple) with the positions on the GDP purine ring labeled by position and atom number. In general, nucleotide binding appears to elicit a contraction in the binding pocket, particularly among three important hydrogen-bonding residues surrounding the nucleobase: Asn119, Asp122, and Asn150.

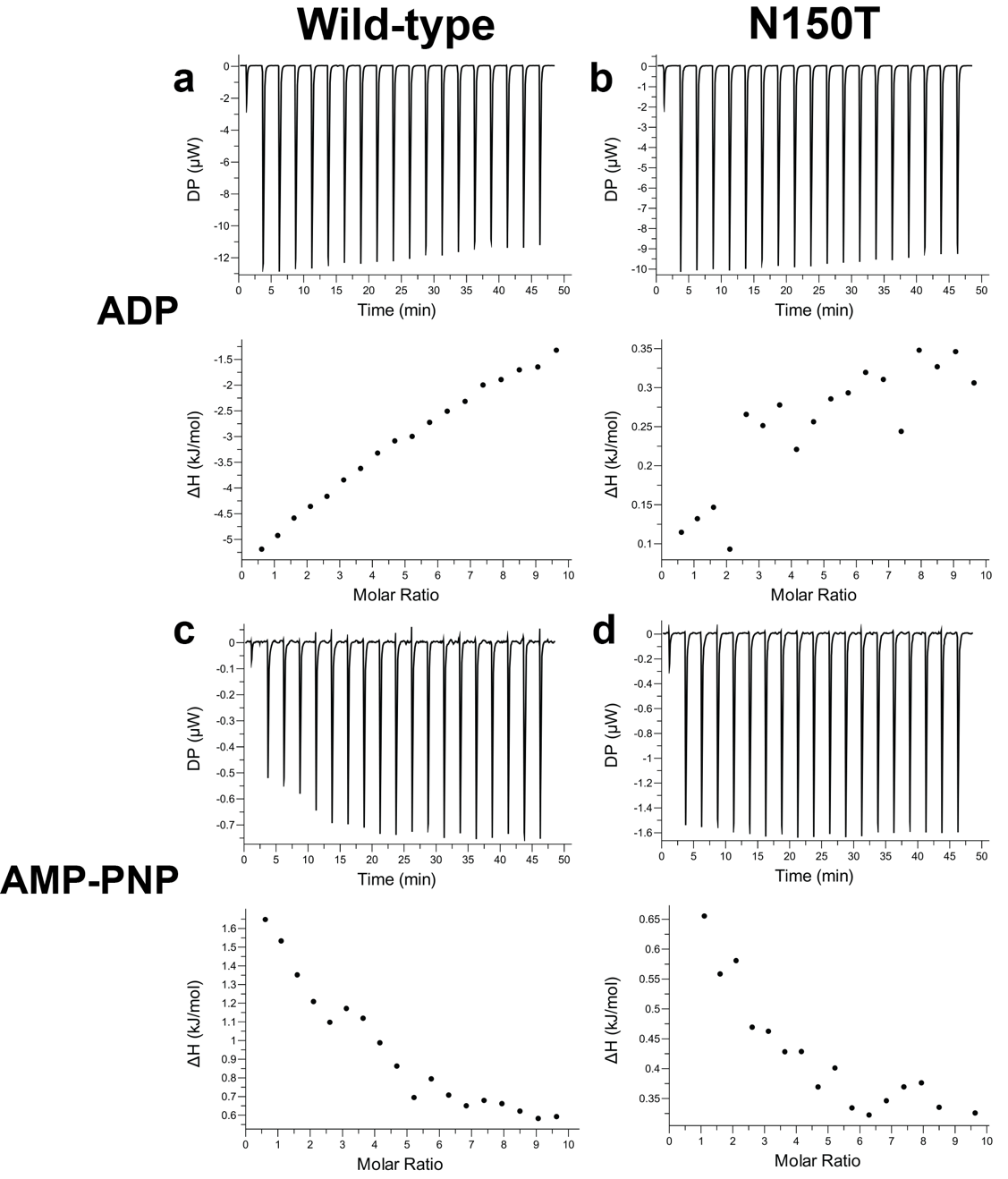

**Fig. S5.** ITC data of adenine-containing nucleotides titrated into *Vc*NFeoB do not indicate stable binding but are most consistent with a high degree of conformational change. Representative ITC thermograms (top) and ΔH vs. molar ratio traces (bottom) of WT (left) and N150T (right) *Vc*NFeoB titrated with either ADP (**a**, **b**) or AMP-PNP (**c**, **d**). All datasets have been corrected for nucleotide dilution into buffer in the absence of protein. Continuous, non-saturating heat evolution suggests weak binding associated with either rapid off rates or with a large conformational change. All values were determined in triplicate and represent the mean ± one standard deviation of the mean.

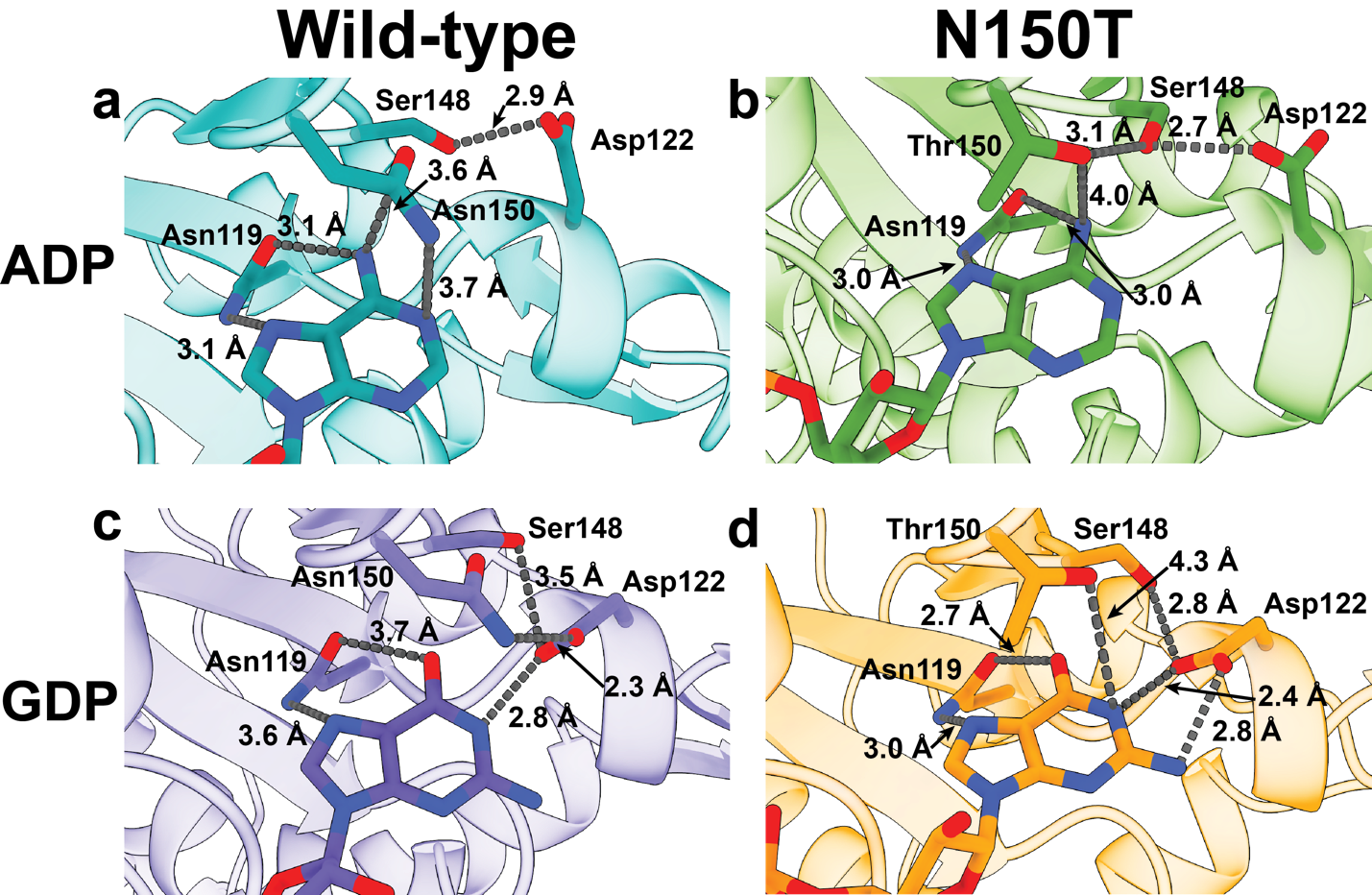

**Fig. S6** AlphaFold3 predicts important structural differences between binding adenine nucleotides and guanine nucleotides to *Vc*NFeoB. Close-up of the nucleotide-binding pocket of WT *Vc*NFeoB (**a**) and N150T *Vc*NFeoB (**b**) in the presence of ADP, based on the AlphaFold3 lowest-energy calculated structures. Close-up of the nucleotide-binding pocket of the ground-truth, experimentally-determined WT *Vc*NFeoB(His)_6_ (**c**) and N150T *Vc*NFeoB(His)_6_ (**d**) structures in the presence of GDP. ADP (and ATP) binding elicits an “Asp off” state in which Asp122 and Ser148 engage in tight hydrogen bonding, opening the nucleotide pocket and weakening overall interactions with the nucleobase, whereas GDP (and GTP) binding elicits an “Asp on” state in which Asp122 actively engages the guanine nucleobase and additional hydrogen bonds are formed within the nucleotide-binding pocket.

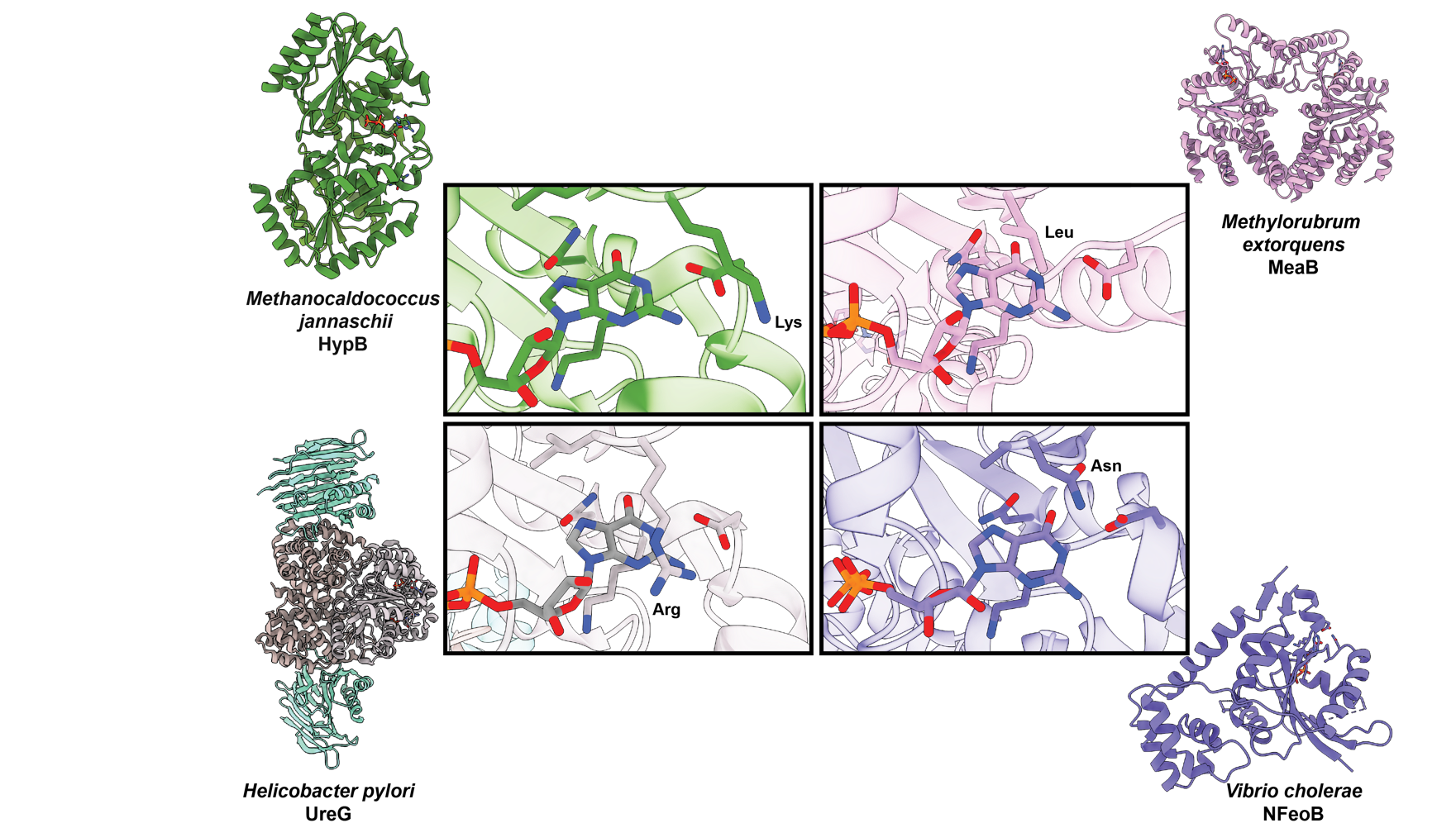

**Fig. S7.** Altered hydrogen bonding interactions about the nucleobase may be a general driver of nucleotide promiscuity among various bacterial NTPases. In particular, variability of the residue analogous to Asn150 in the G5 motif of *Vc*NFeoB (purple) appears to be high in bacterial NTPases, with altered residues in this position including Lys (*Methanocaldococcus jannaschii* HypB, PDB ID 2HF9, green), Leu (*Methylorubrum extorquens* MeaB, PDB ID 2QM7, pink), and Arg (*Helicobacter pylori* UreG, PDB ID 4HI0, pink, gray, and teal). Residues analogous to Asp122 in the G4 motif of *Vc*NFeoB are conserved in all NTPases presented here. We currently hypothesize that flexibility in this region contributes to promiscuity of these types of NTPase enzymes and domains.
